# Intron retention is dynamically regulated during zebrafish larval development and disrupted by high-fat diet

**DOI:** 10.1101/2025.08.18.670806

**Authors:** Jesús Gómez-Montalvo, Jesus Hernandez-Perez, Cecilia Zampedri, Samantha Carrillo-Rosas, Daniela Fernanda Suárez-Bernal, Paula Marroquín-Morales, Marian Farrera-Borraz, Silvia Hinojosa-Alvarez, C. Fabián Flores-Jasso, Rocio Alejandra Chavez-Santoscoy, S. Eréndira Avendaño-Vázquez, Jose Mario Gonzalez-Meljem

**Affiliations:** Tecnologico de Monterrey, School of Engineering and Sciences, Mexico; Consorcio de Metabolismo de RNA y Vesículas Extracelulares, Instituto Nacional de Medicina Genómica, INMEGEN, Mexico City, Mexico

## Abstract

Intron retention (IR) is an understudied mode of alternative splicing with emerging roles in development and stress responses. Here, we present a comprehensive in vivo analysis of IR dynamics during zebrafish larval development and under environmental perturbation. Using deep poly(A)-selected RNA sequencing across three post-fertilization stages (4, 10, and 15 days), we identify IR as a dominant and developmentally regulated splicing event, affecting over 1,000 genes. Unexpectedly, differential IR occurs largely independently of transcript abundance, suggesting a distinct regulatory axis that modulates RNA output via isoform structure rather than expression level. Genes exhibiting IR are enriched in pathways critical to RNA metabolism, neurodevelopment, cell stress responses and genome maintenance—functions that are not captured by differential expression analysis alone. We further show that exposure to a high-fat diet (HFD) reshapes the IR landscape, eliciting both shared and diet-specific splicing responses. Strikingly, some introns display opposing retention dynamics under HFD compared to normal development, highlighting the context-dependent responsiveness of IR regulation. Together, our findings position IR as a selective and environmentally responsive mechanism that contributes to shaping the transcriptome during vertebrate development.

## Introduction

Intron retention (IR) is a form of alternative splicing (AS) in which intronic sequences are retained in mature transcripts. Although initially interpreted as a splicing defect, IR is now understood to function as a regulated mechanism influencing transcript stability, nuclear export, and translational output [1–7]. Compared to other modes of AS—such as skipped exons or alternative splice site usage—IR remains undercharacterized in vertebrates, particularly in vivo. This gap persists despite growing evidence that IR contributes to essential biological processes including differentiation, stress responses, and immune regulation [1,2,8,9].

In contrast to the other modes of AS, IR is distinct in that it often reduces gene expression by preventing translation or promoting transcript degradation, and its regulation has been reported to be uncoupled from changes in transcript abundance [10–14]. This suggests that IR may function as a post-transcriptional layer for modulating gene dosage during development. However, the extent to which IR is developmentally regulated, tissue-specific, or environmentally responsive in vivo remains unclear.

Zebrafish (*Danio rerio*) is a widely used vertebrate model for studying development. Most transcriptomic studies have focused on embryogenesis, while the larval period—which includes major events in tissue maturation and functional differentiation—has received comparatively less attention [15–18]. Limited data suggest that IR occurs in early zebrafish development [19–21], but its prevalence, targets, and temporal behavior during larval stages have not been systematically defined.

To better understand the dynamics of IR during development, we performed poly(A)-selected RNA sequencing on whole zebrafish larvae collected at 4, 10, and 15 days post-fertilization (dpf) to characterize (1) the variation in IR levels across developmental time points; (2) the differential IR across specific gene sets or biological processes; and (3) the responsiveness of IR to environmental stimuli. To explore the latter, we introduced a dietary perturbation— exposure to a high-fat diet (HFD)—as a model of nutritional stress. HFD has been shown in other systems to influence AS, including IR [22,23], and provides a physiologically relevant condition to test the environmental sensitivity of IR regulation during organismal development.

By integrating differential IR, gene expression, and functional enrichment analyses, we show that IR is a widespread and dynamic feature of zebrafish larval development, affecting genes involved in RNA processing, neural and immune functions, and developmental signaling pathways. We further demonstrate that IR patterns represent a pervasive and independent axis of transcriptome regulation—one that modifies RNA output without necessarily altering total gene expression levels.

## Results

### Intron retention emerges as a predominant and regulated splicing feature during zebrafish larval development

To characterize the landscape of AS during zebrafish larval development, we quantified five canonical AS event types—IR, skipped exons (SE), alternative 5′ splice sites (A5SS), alternative 3′ splice sites (A3SS), and mutually exclusive exons (MXE)—across three developmental time points: 4, 10, and 15 dpf.

While all event types were detected, IR and SE predominated at each stage (Fig 1A and S1–S5 Tables). We identified 784 IR events and 893 SE events at 10 vs 4 dpf, and 936 IR events and 1,269 SE events at 15 vs 4 dpf. These proportions contrast with previous observations in zebrafish embryos, where SE accounted for 26–53% of splicing events and IR for ∼8% [19–21]. Here, we found that IR constituted 35–40% of all detected AS events in larval stages, representing a ∼4.7-fold increase in IR frequency and a ∼1.3-fold increase in SE compared to embryonic levels.

**Fig 1.**
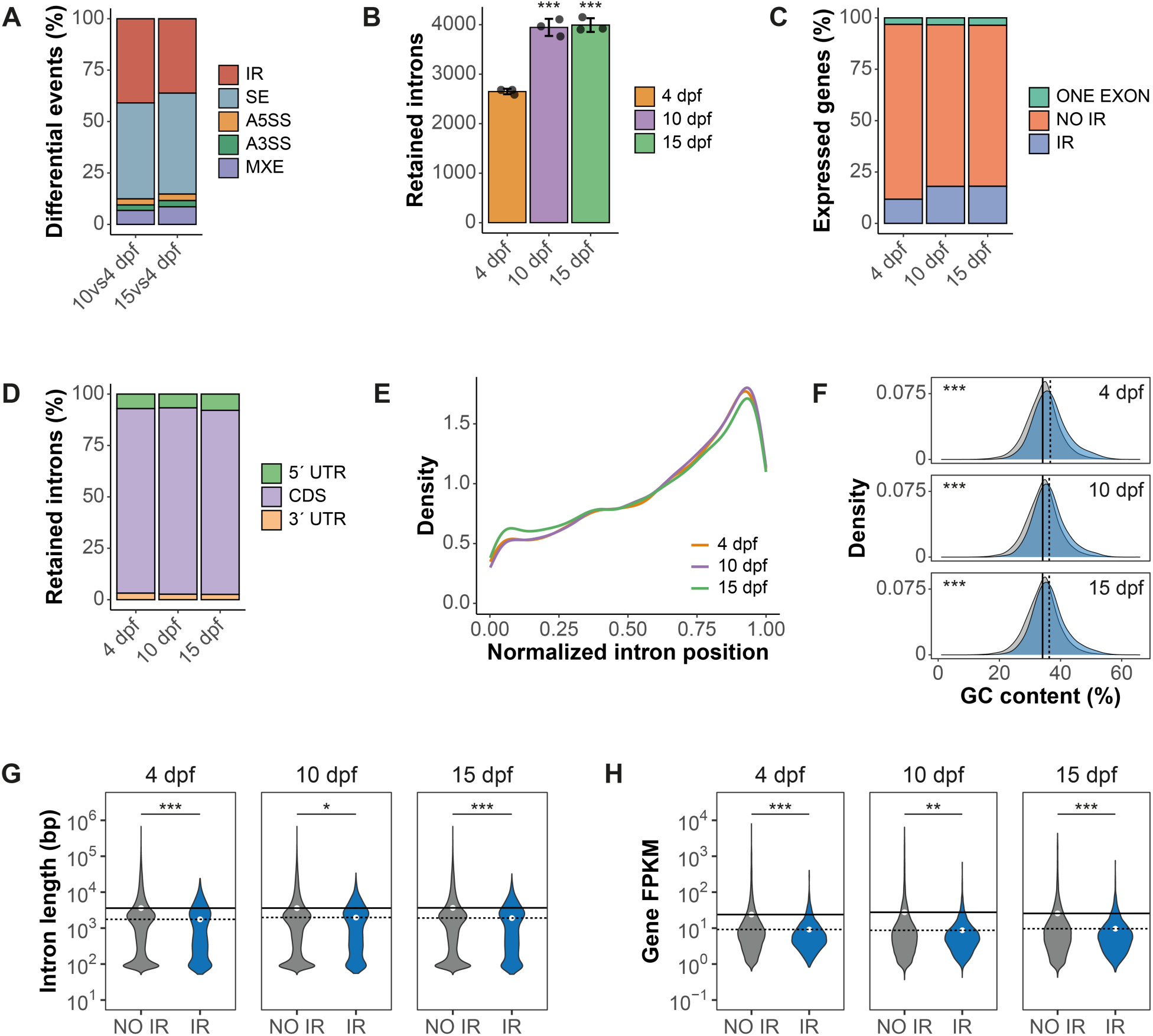
Characterization of retained introns in zebrafish larvae at various developmental time points. **(A)** Proportion of distinct types of AS events during zebrafish larval development. **(B)** Bar plot showing the mean number of retained introns per time point. Black points represent each replicate (n = 3 per time point); error bars denote standard deviation. Asterisks indicate statistically significant differences compared to 4 dpf (one-way ANOVA with Benjamini-Hochberg post-hoc test). **(C)** Proportion of genes with FPKM > 1 categorized as single-exon genes (ONE EXON), no IR (NO IR), or intron-retaining (IR). **(D)** Distributions of retained introns across UTR and coding regions. **(E)** Density plot of normalized intron position within genes (0 = 5′ end; 1 = 3′ end). **(F)** GC content distribution in constitutively spliced (gray) and retained (blue) introns. **(G)** Violin plots showing intron lengths in NO IR and IR categories. **(H)** Transcript levels (FPKM) in genes with or without retained introns. In **(F)–(H)**, solid lines indicate NO IR group mean value, and dotted lines indicate IR group mean value; significance was assessed using the Wilcoxon Rank Sum test (\**p* < 0.01, \*\**p* < 0.001, \*\*\**p* < 0.0001).

The total number of retained introns also increased over time, from a mean of 2,650 ± 54 at 4 dpf to 3,945 ± 172 at 10 dpf and 3,991 ± 138 at 15 dpf (Fig 1B). In parallel, the fraction of genes expressing at least one IR event rose from 12% at 4 dpf to 18% at both 10 and 15 dpf (Fig 1C).

IR events were primarily located within coding regions (CDS) and showed a positional enrichment toward the 3′ end of transcripts (Fig 1D and Fig 1E). Compared to non-retained introns, retained introns had consistently higher GC content (mean GC% at 4 dpf: IR = 36.67%, non-IR = 34.16%; at 10 dpf: IR = 36.27%, non-IR = 34.15%; at 15 dpf: IR = 36.29%, non-IR = 34.13%) (Fig 1F). Intron length also differed, with retained introns being shorter (mean length at 4 dpf: IR = 1,755 bp, non-IR = 3,605 bp; at 10 dpf: IR = 1,972 bp, non-IR = 3,606 bp; at 15 dpf: IR = 1,900 bp, non-IR = 3,661 bp) (Fig 1G). Full annotations are provided in S6 Table.

To examine the relationship between IR and gene expression, we compared transcript abundance between genes with and without retained introns. Across all time points, genes exhibiting IR consistently showed lower expression levels (mean FPKM at 4 dpf: IR = 9.13, non-IR = 23.84; at 10 dpf: IR = 8.71, non-IR = 27.56; at 15 dpf: IR = 9.67, non-IR = 25.64) (Fig 1H).

Together, these findings show that IR is not only widespread during larval development but also exhibits distinct sequence features and associations with reduced transcript abundance, suggesting its role as a regulated component of the developmental transcriptome.

### Intron retention is widespread and largely uncoupled from transcript abundance

To characterize how IR varies during development, we identified differentially retained introns across successive larval stages. Comparisons between 10 and 4 dpf, 15 and 4 dpf, and 15 and 10 dpf yielded 1,382 differential IR events involving 1,068 genes (784 introns at 10 vs 4 dpf; 936 at 15 vs 4 dpf; and 348 at 15 vs 10 dpf) (Fig 2A and S1 Table). Most of these changes reflected increased IR at later stages, consistent with a broader trend observed across the dataset (as shown in Fig 1B).

**Fig 2.**
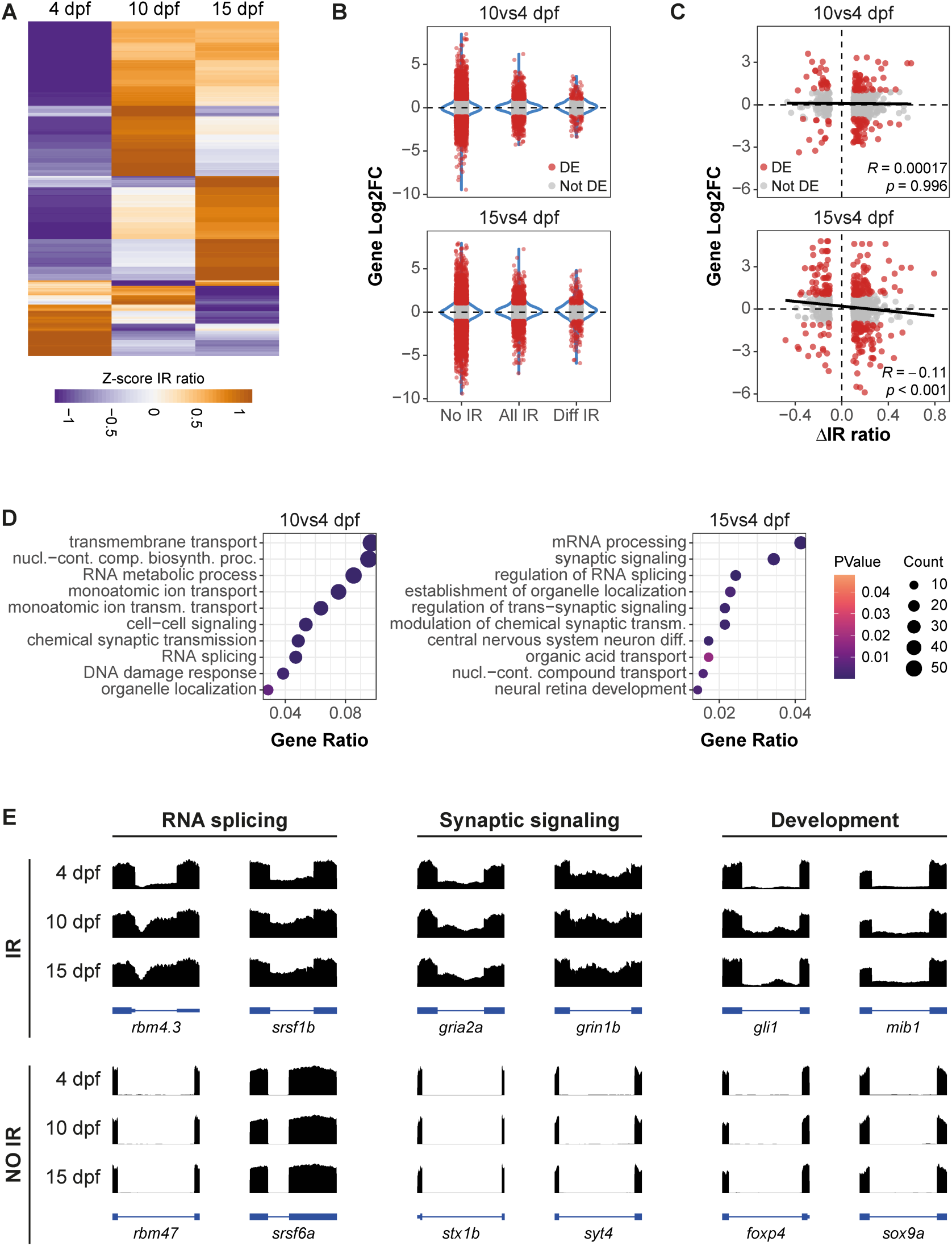
Analysis of intron retention dynamics during zebrafish larval development. **(A)** Heatmap of 1,382 differentially retained introns across 4, 10, and 15 dpf. IR ratio values are Z-score normalized; positive values indicate increased retention. **(B)** Distribution of the Log2 fold change (Log2FC) values for genes with no IR (No IR), any IR (All IR), or differentially retained introns (Diff IR) in the 10 vs 4 dpf and 15 vs 4 dpf contrasts. Each dot represents a gene; differentially expressed (DE) genes are shown in red; violin plots (in blue) depict the distribution of Log2FC values. **(C)** Spearman correlation between changes in IR and transcript abundance for differentially retained introns. Each point represents an intron; red dots indicate cases where the corresponding gene was also DE. Correlation coefficients (*R*) and p-values are indicated for each contrast. **(D)** GO enrichment analysis of genes with differential IR, showing the top 10 enriched biological processes for the 10 vs 4 dpf and 15 vs 4 dpf contrasts. Dot size and color indicate gene count and statistical significance, respectively; Gene Ratio represents the proportion of genes associated with each GO term. **(E)** Representative RNA-seq tracks of genes with differential IR (top) or no IR (bottom) across larval stages. Exons are shown as boxes and introns as lines. Examples are grouped by functional category: RNA splicing (IR: *rbm4.3*, *srsf1b*; NO IR: *rbm47*, *srsf6a*), synaptic signaling (IR: *gria2a*, *grin1b*; NO IR: *stx1b*, *syt4*), and development (IR: *gli1*, *mib1*; NO IR: *foxp4*, *sox9a*). Intron IDs are provided in S2 Fig.

To determine whether IR regulation is accompanied by changes in transcript abundance, we quantified differentially expressed (DE) genes across the same developmental intervals. In total, 9,809 genes were DE in at least one comparison (5,930 at 10 vs 4 dpf; 8,047 at 15 vs 4 dpf; 2,371 at 15 vs 10 dpf), covering roughly 30% of annotated genes in zebrafish (Fig S1A and S7 Table).

We then classified expressed genes (FPKM > 1) into three categories: those with no detectable IR (No IR), those with IR events regardless of change (All IR), and those with differential IR (Diff IR) (Fig 2B). In the 10 vs 4 dpf comparison, 23% of No IR genes were also DE, compared to 17% and 18% in the All IR and Diff IR groups, respectively. At 15 vs 4 dpf, 31% of No IR genes were DE, versus 25% in the All IR group and 30% in the Diff IR group.

The range of log2 fold change (Log2FC) in expression also differed among categories. Genes without IR exhibited the broadest expression variability (−9.47 to 8.49 at 10 vs 4 dpf; −9.4 to 7.94 at 15 vs 4 dpf), while genes with differential IR showed a narrower Log2FC range (−3.36 to 3.61 and −5.89 to 4.8, respectively) (Fig 2B). Most genes with differential IR were not differentially expressed—82% in the 10 vs 4 dpf contrast and 70% in the 15 vs 4 dpf contrast (Fig 2B and Fig 2C).

We also examined the relationship between IR levels and transcript abundance. No correlation was observed in the 10 vs 4 dpf comparison (Spearman *R* = 0.00017), and only a weak inverse correlation at 15 vs 4 dpf (*R* = −0.11) (Fig 2C). These findings indicate that developmental changes in IR are often uncoupled from transcript abundance regulation.

Together with the observation that most differentially retained introns occur in genes without differential expression, these results suggest a broader regulatory principle: that IR modifies RNA output through changes in splicing rather than transcript quantity. Hence, our findings position IR as a pervasive and independent axis of transcriptome regulation during development—one that remodels RNA output without necessarily altering gene expression levels.

### Intron retention targets gene networks central to RNA processing, neurodevelopment, and genome maintenance

The observation that IR often occurs independently of changes in transcript abundance raises the question of which biological functions are regulated through this mechanism. To address this, we examined the identity of genes exhibiting differential IR across development.

We first compared pathway and gene ontology (GO) enrichment patterns between differentially expressed (DE) genes and those with differential intron retention (Diff IR). DE genes showed enrichment for gene sets and ontology terms related to immune signaling, cell cycle control, oxidative phosphorylation, and fatty acid metabolism (Fig S1B and S8–S9 Tables). In contrast, Diff IR genes revealed enrichment for biological processes involving RNA splicing, synaptic function, and neuronal development (Fig 2D and S10 Table).

Genes involved in RNA metabolism were prominently represented among IR targets, including splicing regulators from the RBM and SRSF families (Figs 2D, 2E and S2). Consistent with this, IR was often detected in transcripts encoding spliceosomal components, suggesting a layer of autoregulation among the splicing machinery.

IR was also enriched among genes associated with synaptic signaling and transmembrane transport. These included glutamatergic synapse components such as *gria2a* and *grin1b*, which accounted for approximately 30% of the genes in synaptic signaling categories (Fig 2E and S2 Fig). Notably, genes related to neurodevelopment showed dynamic IR, including *nkx3-2*, *hmga1b*, *gli1*, *numb*, and *mib1*. In *gli1*, IR rose from 0.07 at 4 dpf to 0.38 at 10 dpf before declining to 0.15 at 15 dpf, while in *numb* and *mib1*, IR increased steadily over time. In *dlc*, distinct introns exhibited opposite retention trajectories, suggesting intron-level specificity in splicing regulation (S2 Fig).

Processes related to central nervous system differentiation and neural retina development were also overrepresented among Diff IR genes (Fig 2D), aligning with the timing of neuronal maturation during this period [24]. These findings indicate that IR affects genes involved in both the general splicing program and tissue-specific developmental functions, particularly in the nervous system.

In addition to neuronal and RNA-processing genes, we identified differential IR in genes involved in DNA repair. Enrichment of DNA damage response terms included core components of the Fanconi anemia pathway (*fan1*, *fance*, *fancg*) and nucleotide excision repair (*ercc5*, *ercc8*), pointing to potential roles for IR in genome maintenance during larval growth (Fig 2D and S2 Fig).

Although immune- and lipid-related terms were not significantly enriched in global analyses, manual inspection revealed IR in several genes of the complement cascade (*c4b*, *c6*, *c7a*, *c8b*) and lipid transport or metabolism (*lipea*, *lipeb*, *ldlrap1a*, *lrp8*, *hdlbpb*), suggesting more targeted effects in these pathways (S2 Fig).

In many cases, IR changes occurred in the absence of differential gene expression (S2 Fig), reinforcing its role as a post-transcriptional mechanism modulating RNA output across developmental transitions. Our GO analysis supports the concept that IR is dynamically regulated across development and selectively enriched in gene networks involved in RNA processing, neurodevelopment, DNA repair, and metabolic homeostasis.

### Dietary perturbation reshapes intron retention dynamics during early development in zebrafish

Having established that IR is developmentally regulated and often decoupled from changes in transcript abundance, we next asked whether IR is sensitive to environmental conditions. To test this, we used dietary intervention as a model of physiological stress, focusing on a high-fat diet (HFD), which alters metabolic signaling in zebrafish and other vertebrates [25].

Larvae were exposed to HFD from 6 to 10 dpf, resulting in visible lipid accumulation in the liver and intestinal tract by 10 dpf (S3A Fig). We refer to these HFD-fed larvae as “10HFD” and to their age-matched standard-fed controls as “10 dpf.” IR dynamics were analyzed in three contrasts: normal development (10 vs 4 dpf), HFD development (10HFD vs 4 dpf), and diet change (10HFD vs 10 dpf). The first comparison was already described in previous sections.

IR and skipped exons (SE) remained the predominant splicing modes under both HFD and standard conditions (Fig 3A and S1–S5 Tables). We identified 524 IR and 602 SE events in HFD development, and 205 IR and 228 SE events in the diet change contrast, indicating that both development and diet influence splicing patterns.

**Fig 3.**
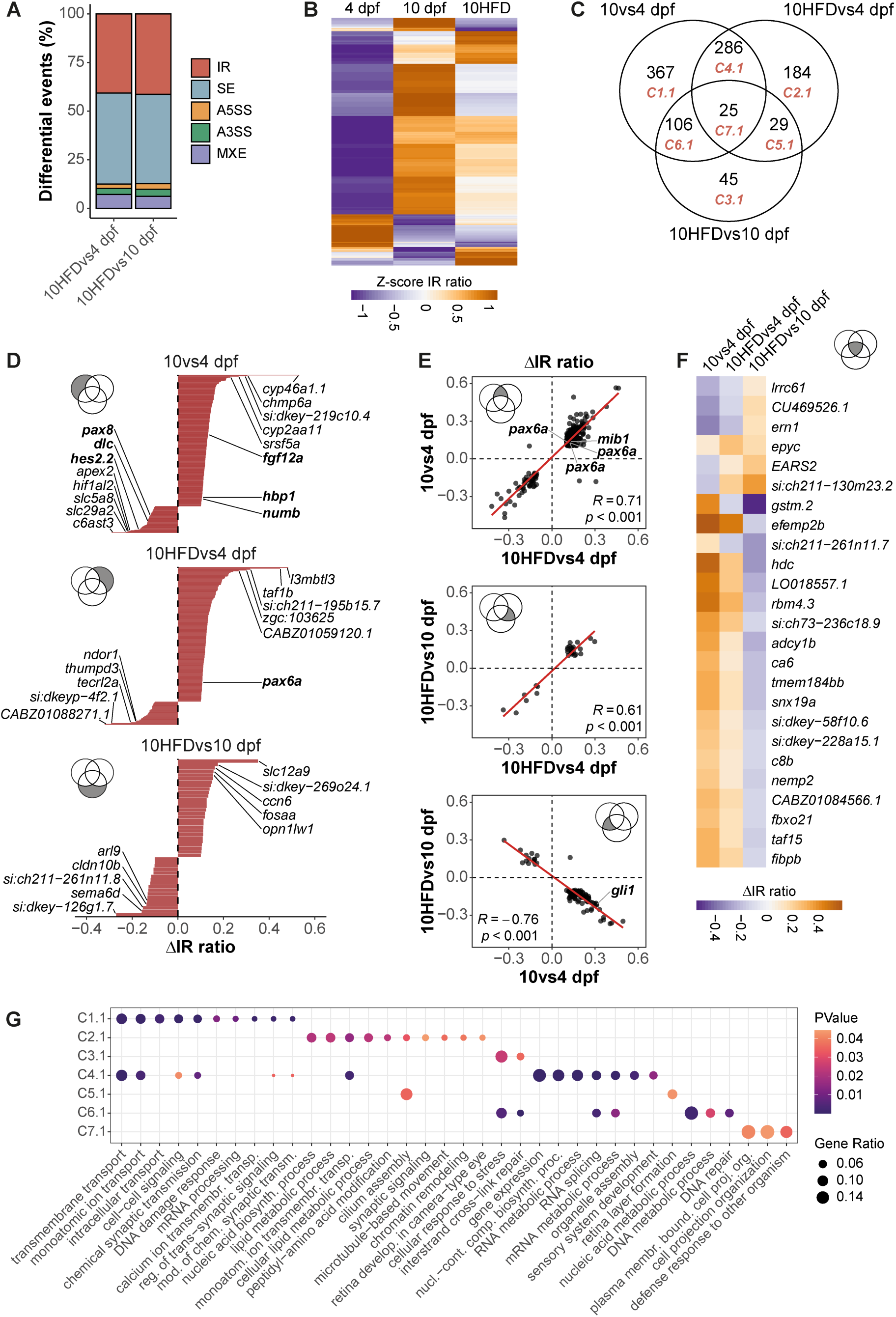
Differential IR analysis in zebrafish larvae fed with HFD. **(A)** Proportion of AS event types identified in the HFD development (10HFD vs 4 dpf) and diet change (10HFD vs 10 dpf) contrasts. **(B)** Heatmap showing Z-score normalized IR ratios for 1,042 differentially retained introns across 4 dpf, 10 dpf, and 10HFD samples. Differential events were identified in the 10 vs 4 dpf, 10HFD vs 4 dpf, and 10HFD vs 10 dpf contrasts. Positive and negative Z-scores reflect increased and decreased IR, respectively. **(C)** Venn diagram summarizing the overlap of differentially retained introns across the three contrasts. **(D)** Bar plots depicting changes in IR levels for introns uniquely altered in each contrast. For each group, the five most strongly upregulated and downregulated introns are shown; developmental genes are indicated in bold. **(E)** Spearman correlation analyses of IR changes for introns shared between contrasts. The correlation coefficient (*R*) and p-value are reported for each comparison. Genes of developmental relevance are highlighted in bold. **(F)** Heatmap of IR changes for the 25 introns differentially retained across all three contrasts. In **(D–F)**, the corresponding subset of the Venn diagram in **(C)** is indicated. **(G)** Dot plot showing significantly enriched GO biological process terms for each cluster of differentially retained introns in **(C)**. Gene Ratio represents the proportion of intron-retaining genes associated with each GO term.

We next assessed whether HFD modifies IR profiles. Compared to standard-fed controls at 10 dpf, HFD-fed larvae exhibited distinct IR changes (Fig 3B). Of the differentially retained introns, 367 were specific to normal development, 184 were unique to HFD development, and 45 occurred only in response to the dietary shift (10HFD vs 10 dpf) (Fig 3C, Fig 3D and S11 Table). Notably, 82% of these introns showed increased retention, consistent with the broader trend observed during larval development.

To further explore the interaction between diet and developmental regulation, we examined overlap and directionality across contrasts. A subset of 286 introns changed retention in both standard and HFD development, with strongly correlated IR ratios (*R* = 0.71) (Fig 3E, top). A smaller group of 29 introns exhibited altered retention in response to HFD regardless of developmental stage, also showing consistent directional changes (*R* = 0.61) (Fig 3E, middle).

Unexpectedly, we identified 106 introns with opposing regulation between normal development and diet change comparisons, marked by a negative correlation in ΔIR ratios (*R* = – 0.76) (Fig 3E, bottom). These introns increased retention during development under standard diet but showed reduced retention when larvae were exposed to HFD. For example, *gli1* displayed increased IR during development (ΔIR = +0.31) but decreased IR under HFD (ΔIR = –0.22). Similar patterns were observed in 25 introns that were shared across all three contrasts, reinforcing the context-dependent regulation of IR (Fig 3F).

These findings indicate that a high-fat diet alters the developmental IR landscape in zebrafish larvae. Subsets of introns exhibit specific or even inverse regulation in response to diet, revealing that IR is not only developmentally programmed but also responsive to environmental inputs during early growth.

### High-fat diet selectively alters IR in genes linked to stress response, RNA metabolism, and developmental signaling

Next, we further characterized the biological processes affected by IR under HFD conditions. We analyzed the functional categories associated with Diff IR across contrasts. Groups of Diff IR genes were defined based on their presence in the intersections of the three pairwise comparisons (Fig 3G and S12 Table).

Genes with IR changes exclusive to normal development were enriched in processes such as transmembrane and ion transport, chemical synaptic transmission, mRNA processing, and DNA damage response. Manual inspection of this group revealed multiple developmental regulators with dynamic IR patterns, including *hbp1*, *numb*, *dlc*, and *hes2.2* (Fig 3D, top).

Genes with IR changes specific to HFD development were enriched for nucleic acid biosynthesis, lipid metabolism, and synaptic signaling. In contrast, genes affected only by the diet change (10HFD vs 10 dpf) showed enrichment for cellular stress response and interstrand cross-link repair, suggesting acute adaptation to dietary perturbation.

The subset of genes with shared IR changes across normal and HFD development included those involved in RNA processing, gene expression, synaptic signaling, and sensory system development. Genes overlapping between the HFD development and diet change contrasts showed more limited enrichment, with terms related to cilium assembly and retina layer formation.

Notably, the group of genes with inverse IR responses between normal development and diet change showed enrichment for nucleotide metabolism, DNA repair, RNA processing, and stress response—highlighting potential regulatory conflicts between endogenous developmental programs and exogenous dietary input. Finally, genes with IR changes common to all three contrasts were associated with cell projection organization and immune defense processes.

A comparison with the DE gene set revealed minimal overlap in enriched biological terms. Of all GO terms identified across both datasets, only ∼20% were shared. While DE genes— particularly those altered in both HFD development and diet change—showed consistent enrichment for lipid-related pathways (e.g., sterol, steroid, and cholesterol biosynthesis and transport), lipid metabolism was only marginally represented among Diff IR gene annotations (S3B–S3D Figs and S12–S13 Tables).

Taken together, our analysis shows that dietary fat modifies the IR landscape across a broad range of biological functions, many of which extend beyond canonical lipid metabolic pathways. The limited overlap between IR- and expression-based enrichments further supports the view that IR operates in a distinct regulatory layer during environmental and developmental transitions.

## Discussion

This study reveals that IR is a widespread and developmentally dynamic component of the zebrafish transcriptome. By profiling zebrafish larvae at three post-fertilization stages and under dietary perturbation, we demonstrate that IR affects a significant number of genes across key biological processes—most notably RNA processing, developmental programs, stress responses, and DNA repair.

One of the central findings of this work is that IR regulation often occurs independently of transcript abundance. The majority of differentially retained introns arose in genes with stable expression levels, and IR magnitude showed little to no correlation with transcript abundance. While previous studies have reported inverse relationships between IR and transcript levels, many of these examples involve Diff IR genes exhibiting only modest expression changes (|Log2FC| < 1) [8,26]. Likewise, studies examining IR dynamics across time during aging have similarly found that Diff IR genes generally maintain stable transcript levels [13]. Hence, IR emerges as a widespread and distinct regulatory axis within the transcriptome—one that reconfigures RNA structure independently of changes in overall gene expression levels.

Although alternative splicing coupled to nonsense-mediated mRNA decay (AS-NMD) has been proposed as a regulatory mechanism in vertebrates [27,28], our data do not necessarily support this as a general mode in zebrafish larvae. Genes with differential IR were generally not differentially expressed and displayed narrower Log2FC distributions than genes lacking IR, suggesting that IR does not broadly act as a trigger for transcript downregulation in this context. One plausible explanation is that coupling between IR and NMD may require specific splicing regulators such as SRSF1, known to promote exon junction complex deposition and enhance NMD efficiency [29]. The apparent lack of IR transcript destabilization in our dataset may thus reflect limited activity of such coupling factors during larval development, supporting the view that IR primarily modulates RNA output through isoform diversity rather than decay.

This distinction challenges a common assumption in transcriptome studies that expression changes are the primary indicator of gene regulation. Our functional enrichment analyses show minimal overlap between DE and diff IR gene sets. For example, while DE genes were enriched in canonical metabolic pathways such as lipid biosynthesis and transport, IR-affected genes were more strongly associated with RNA metabolism, synaptic signaling, and genome maintenance. This decoupling has also been observed by other groups [3,30] and reinforces the notion that IR acts as a post-transcriptional layer of regulation, complementary to but distinct from transcript abundance.

Importantly, our results suggest that earlier transcriptome studies may have underestimated the prevalence of IR during development [19–21]. This may reflect technical limitations: prior analyses typically used lower sequencing depth (<10 million reads) and splicing tools designed for exon skipping or splice site changes. In contrast, our study employed high-depth RNA-seq (averaging 150 million reads) and IRFinder [14], a tool tailored to quantifying retained introns. These methodological advances likely improved sensitivity, revealing regulatory features that would otherwise be missed. Thus, our findings also underscore the value of specialized computational approaches in decoding post-transcriptional complexity.

The environmental sensitivity and specificity of IR adds further evidence supporting its regulatory role. We show that HFD exposure alters the IR landscape across hundreds of genes, including those involved in synaptic function, cellular stress, and DNA repair. Strikingly, many IR changes observed under HFD were specific, and in some cases, inverse to those seen in normal development—highlighting that IR regulation is both developmentally programmed and environmentally responsive. However, only a minority of these HFD-associated IR genes were directly related to lipid metabolism, implying that nutritional stress affects a broad range of transcriptomic functions beyond canonical metabolic pathways. This view is further supported by studies showing that tissues from rodents fed a high-fat diet displayed altered AS events— including IR—primarily in non-lipid-related genes [22,31].

Together, these findings position IR as a selective and responsive mechanism that modulates transcriptome structure without necessarily altering RNA quantity. While IR has been described in multiple systems, our study provides an organism-wide in vivo demonstration of its prevalence, developmental regulation, and environmental responsiveness. By focusing on IR specifically, and with sufficient sequencing depth, we provide evidence that it is not a marginal or passive event, but a regulated component of transcriptome architecture.

Building on previous observations from our group and others showing that age-related IR changes occur largely independently of transcript abundance [13,32], this study expands this paradigm to vertebrate development. We thus propose that IR represents a functionally distinct mode of transcriptome regulation—one that governs RNA fate rather than mere RNA presence. Retained introns may alter subcellular localization, impede translation, or modulate a transcript’s interaction with regulatory proteins and other non-coding RNAs, ultimately determining its stability and functional lifespan [14,33–38]. By influencing RNA fate, we postulate that IR adds a regulatory layer that is complementary to transcriptional output, enabling cells to fine-tune specific gene function during development and in response to physiological cues without altering overall transcript levels.

## Materials and Methods

### Zebrafish

Wild-type zebrafish (*Danio rerio*, AB strain) were bred from natural matings and raised at 28.5 °C under standard facility conditions. At 5 days post-fertilization (dpf), larvae were fed a standard commercial diet (Azoo Ultra Fresh Micron, 0.25% w/v). At 6 dpf, larvae were randomly assigned to two groups: one remained on the standard diet until 10 or 15 dpf; the other received a high-fat diet (HFD), prepared by supplementing the standard diet with boiled egg yolk (2.5% w/v), until 10 dpf. Each sample comprised 20 larvae. Larvae were euthanized by triple PBS wash, immersion in ice-cold water (4 °C, 1 min, until swimming ceased) [39], and immediate transfer to TRIzol reagent. All procedures complied with institutional animal care guidelines and international regulations (EU Directive 2010/63, CCAC, and relevant U.S. standards).

### Nile Red Staining

Lipid accumulation was assessed using Nile Red staining. A stock solution (1 mg/mL) was prepared in acetone. Larvae were incubated for 30 minutes at 28.5 °C in 10 mL aquarium water containing 1 µL of the stock solution. Following staining, larvae were anesthetized with tricaine (0.4% in aquarium water) and embedded in 1% low-melting-point agarose on concave microscope slides. Fluorescent images were acquired for 7–8 larvae per group using an epifluorescence microscope.

### RNA Extraction

Total RNA was extracted from zebrafish larvae homogenized in 500 µL TRIzol using a motorized pestle (Argos Technologies, cat. no. 3-33701). After 5 min incubation at room temperature, 100 µL chloroform were added, followed by a 3 min incubation and centrifugation at 12,000 × g for 15 min at 4 °C. The aqueous phase (200 µL) was mixed with 170 µL isopropanol, incubated for 10 min, and centrifuged at 12,000 × g for 10 min at 4 °C. Pellets were washed with 1 mL of 75% ethanol and centrifuged at 7,500 × g for 5 min at 4 °C. RNA was resuspended in 30 µL RNase-free water. Concentration and purity were measured with an NP80 Implen NanoPhotometer; integrity was assessed by agarose gel electrophoresis.

### RNA Sequencing

RNA integrity was assessed using the Agilent 2100 Bioanalyzer. Poly(A) RNA selection and library preparation were performed with the TruSeq Stranded mRNA kit (Illumina) according to the manufacturer’s protocol. Library concentration and fragment size distribution were evaluated using the Qubit Fluorometer and Agilent TapeStation 4200, respectively. Sequencing was performed on a NovaSeq 6000 S4 flow cell across four lanes, generating ∼200 million paired-end reads (2 × 110 bp) per sample at the National Genomic Sequencing Laboratory Tec-BASE (Tecnológico de Monterrey).

### Data Processing and Quality Control

The quality of raw RNA-seq reads was assessed using FastQC v0.12.1. Sequencing duplicates were identified and removed using the Clumpify tool from the BBTools suite v39.06. Adapter sequences and low-quality bases (Phred score < 30) were trimmed using Cutadapt v4.6. Filtered reads were aligned to the *Danio rerio* reference genome (GRCz11) using STAR v2.7.11b [40], with genome indices generated from ENSEMBL gene annotation v102. Quality summaries from trimming and mapping were compiled using MultiQC v1.20 [41]. After filtering, an average of approximately 150 million paired-end reads per sample were retained, with approximately 80% mapping uniquely to the genome (S4A Fig).

Gene body coverage was evaluated using the *geneBody_coverage.py* function from the RSeQC package v5.0.3 [42], confirming the absence of 3′ bias in the sequencing libraries (S4B Fig). Gene-level counts were computed using featureCounts v2.0.6 [43] with ENSEMBL annotation v102. Count normalization and principal component analysis (PCA) were performed using DESeq2 v1.40.2 [44]. PCA based on the top 500 most variable genes showed tight clustering of biological replicates and clear segregation by developmental time point (S4C Fig).

### Intron Retention Analysis

Retained introns were identified using IRFinder v1.3.1 [14], with references built from the *Danio rerio* ENSEMBL annotation v102. IRFinder quantifies intron retention using the IR ratio, defined as the proportion of transcripts retaining a given intron. Only introns not overlapping other annotated genomic features were analyzed. A total of 219,347 introns from 23,613 genes were assessed.

An intron was considered retained if it met all of the following thresholds: Coverage ≥ 0.8, IntronDepth ≥ 5, SpliceExact ≥ 5, and IRratio ≥ 0.1. These criteria excluded low-confidence events and ensured that retained introns represented ≥10% of gene transcripts with detectable expression (FPKM > 1). Differential IR analysis was conducted using the Likelihood Ratio Test in DESeq2 v1.40.2 [44]. Introns with |ΔIR ratio| > 0.1 and adjusted p-value (padj) < 0.05 were considered differentially retained. RNA-seq coverage tracks for intron-retaining genes were generated using SparK [45].

### Analysis of the Location and Position of Retained Introns

Transcript coordinates for 5′ and 3′ untranslated regions (UTRs) were obtained using the *fiveUTRsByTranscript* and *threeUTRsByTranscript* functions from the GenomicFeatures package v1.52.1. Retained introns located within a UTR of any transcript isoform were classified as UTR introns; all others were categorized as coding sequence (CDS) introns.

Relative intron position within genes was calculated by subtracting the gene start coordinate from the intron end coordinate and dividing the result by the total gene length. Values approaching 0 denote proximity to the 5′ end; values near 1 indicate the 3′ end. Analyses were performed using the ENSEMBL zebrafish annotation v102.

### Analysis of Intron GC Content

Introns were categorized as retained or non-retained. Non-retained introns were defined as those with IntronDepth < 5 and IR ratio < 0.01, following established criteria [46]. Intron sequences were extracted using the *getfasta* function from BEDtools v2.27.1 [47], and GC content was calculated with the *geecee* function from the EMBOSS package v6.6.0.0.

### Alternative Splicing Analysis

Alternative splicing (AS) events excluding IR—including skipped exons (SE), alternative 5′ splice sites (A5SS), alternative 3′ splice sites (A3SS), and mutually exclusive exons (MXE)—were identified using rMATS v4.1.2 [48]. AS events were filtered using maser v1.18.0 to retain those supported by ≥10 average reads. Differential AS events were defined by |ΔPSI| > 0.1 and False Discovery Rate (FDR) < 0.05. Only events in genes with FPKM > 1 were included in downstream analyses.

### Differential Gene Expression Analysis

Differential gene expression analysis was conducted using DESeq2 v1.40.2 [44]. Fold change shrinkage was applied using the ashr method to improve effect size estimation. Genes were defined as differentially expressed if they satisfied both criteria: |Log2 fold change| > 1 and padj < 0.05. The padj values were computed using the Benjamini-Hochberg method for multiple testing correction.

### Gene Set Enrichment Analysis

Differential gene expression lists were ranked by the Wald statistic in descending order. Pre-ranked gene set enrichment analysis (GSEA) was performed using GSEA tool v4.2.2 [49] with default parameters. Hallmark and KEGG gene sets were sourced from the Molecular Signatures Database. Gene sets with p-value < 0.05 and FDR < 0.25 were considered significantly enriched.

### Gene Ontology Analysis

Gene ontology (GO) enrichment analysis was conducted using DAVID (https://david.ncifcrf.gov/) [50], focusing on the GOTERM_BP_ALL category. Redundant GO terms were reduced using REVIGO [51]. For intron-retaining genes, significant enrichment was defined by p-value < 0.05 and fold enrichment > 2. For differentially expressed (DE) genes, GO terms with Benjamini-Hochberg padj < 0.05 were considered significant.

## Data Availability

Raw RNA-seq data have been deposited in the Gene Expression Omnibus (GEO) under accession number **GSE290628**. Analysis code is available from the corresponding author upon request.

## Ethics statement

All experiments conducted on zebrafish larvae were previously approved by the Institutional Committee for Care and Use of Laboratory Animals at Tecnologico de Monterrey (protocol ID 2025-016).

## Acknowledgments

We thank the members of the Senescence and RNA Metabolism Laboratories for their valuable feedback and insightful discussions. We also acknowledge the Instituto Nacional de Medicina Genómica (INMEGEN) for providing computational resources. Library preparation was performed at the Sequencing Unit (INMEGEN) and sequencing was carried out at the National Genomic Sequencing Laboratory Tec-BASE (Tecnologico de Monterrey). We are grateful to MD Valentín Mendoza Rodríguez and QFB Cristina Aranda Fraustro for their technical support.

## Funding disclosure

CFFJ and SEAV are supported by grant numbers A1-S-38213 and 289862 from Secretaría de Ciencia, Humanidades, Tecnología e Innovación (SECIHTI); URL: https://secihti.mx/ and grant numbers 09/2019/I, 08/2017/I, and 08/2019/I from Instituto Nacional de Medicina Genómica (INMEGEN); URL: https://www.inmegen.gob.mx/. JMGM was supported by the “2022 Core Lab Genomics Tec-BASE Seed Fund for Research Projects” from Tecnologico de Monterrey; URL: https://tec.mx/es/investigacion. JGM is supported by a SECIHTI scholarship (817529, CVU 1049823). Funders did not play any role in study design, data collection and analysis, decision to publish or preparation of the manuscript.

## Competing interests

The authors declare that no competing interests exist.

## Related Manuscripts

The authors declare they do not have a related or duplicate manuscript under consideration (or accepted) for publication elsewhere.

## Author contributions

**Conceptualization:** S. Eréndira Avendaño-Vázquez, José Mario González-Meljem

**Data curation:** Jesús Gómez-Montalvo

**Formal analysis:** Jesús Gómez-Montalvo, S. Eréndira Avendaño-Vázquez, José Mario González-Meljem

**Funding acquisition:** C. Fabián Flores-Jasso, Rocio Alejandra Chavez-Santoscoy, S. Eréndira Avendaño-Vázquez, José Mario González-Meljem

**Investigation:** Jesús Gómez-Montalvo, S. Eréndira Avendaño-Vázquez, Jose Mario González-Meljem

**Methodology:** Jesús Gómez-Montalvo, Jesús Hernández-Pérez, Cecilia Zampedri, Samantha Carrillo-Rosas, Daniela Fernanda Suárez-Bernal, Paula Marroquín-Morales, Marian Farrera-Borraz, Silvia Hinojosa-Álvarez

**Project administration:** S. Eréndira Avendaño-Vázquez, Jose Mario Gonzalez-Meljem

**Resources:** Cecilia Zampedri, Samantha Carrillo-Rosas, C. Fabián Flores-Jasso, Rocio Alejandra Chavez-Santoscoy, S. Eréndira Avendaño-Vázquez, Jose Mario Gonzalez-Meljem

**Software:** Jesús Gómez-Montalvo, Silvia Hinojosa-Álvarez

**Supervision:** S. Eréndira Avendaño-Vázquez, José Mario González-Meljem

**Visualization:** Jesús Gómez-Montalvo, S. Eréndira Avendaño-Vázquez, José Mario González-Meljem

**Writing – original draft:** Jesús Gómez-Montalvo, José Mario González-Meljem

**Writing – review & editing:** Jesús Gómez-Montalvo, C. Fabián Flores-Jasso, S. Eréndira Avendaño-Vázquez, José Mario González-Meljem

## Supporting information

### Supplemental Figures

**S1 Fig.**
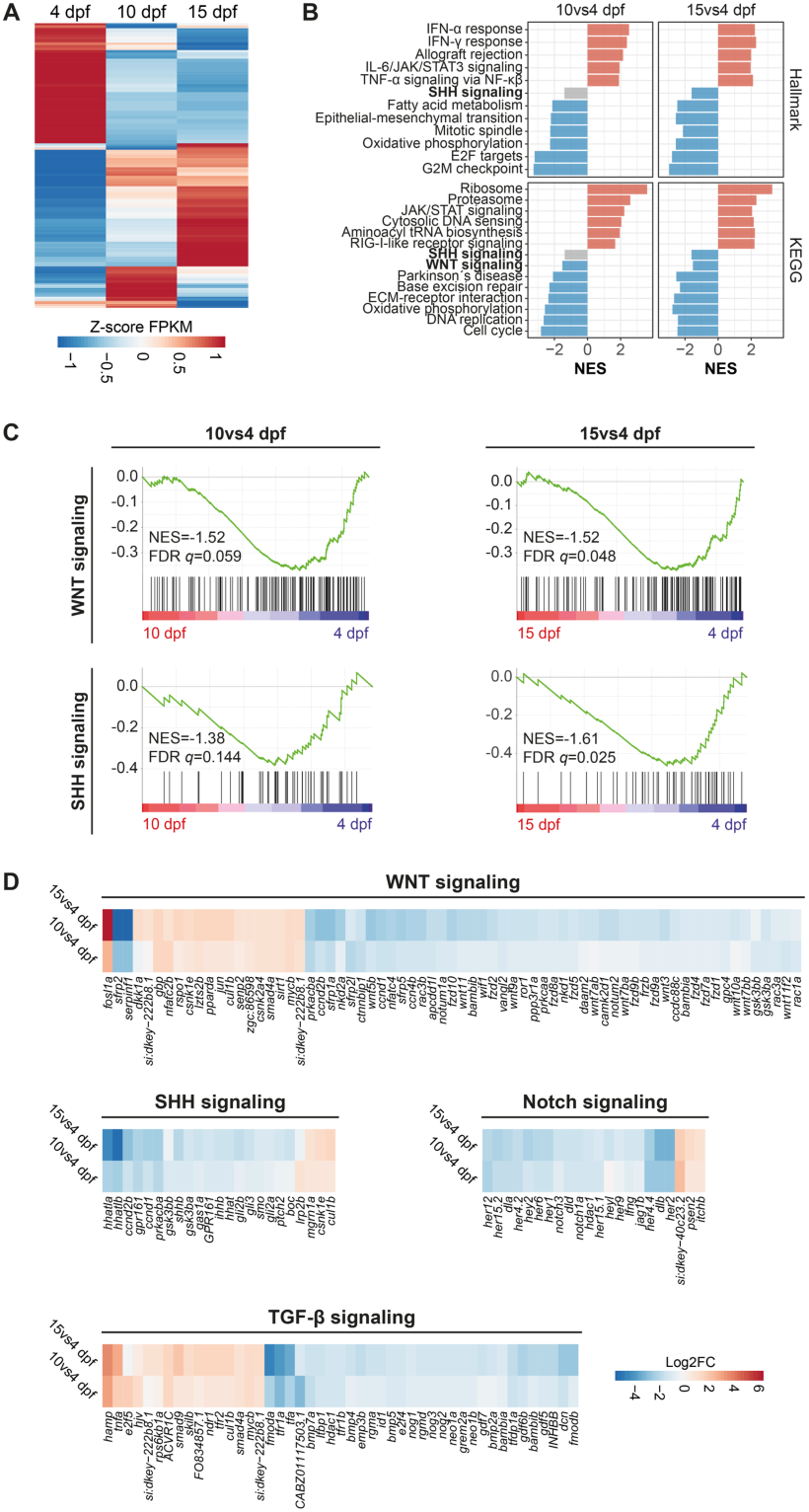
Transcriptomic changes during zebrafish larval development. **(A)** Heatmap of 9,809 differentially expressed (DE) genes across 4, 10, and 15 dpf. Values are Z-score–normalized transcript levels. **(B)** Bar plot of Hallmark gene-set enrichment (GSEA). Top five significantly up-regulated (red) and down-regulated (blue) pathways per developmental contrast are shown; non-significant sets in grey. Development-related pathways are bolded. NES, normalized enrichment score. **(C)** GSEA enrichment plots for KEGG WNT and SHH signaling pathways, both enriched among down-regulated genes. Green curve indicates enrichment score across the ranked gene list (red = up-regulated, blue = down-regulated). Right-skewed enrichment denotes pathway down-regulation. **(D)** Heatmaps of log2 fold change (Log2FC) for DE genes in WNT, SHH, Notch, and TGF-β signaling (|Log2FC| > 1; padj < 0.05). Gene sets curated from KEGG.

**S2 Fig.**
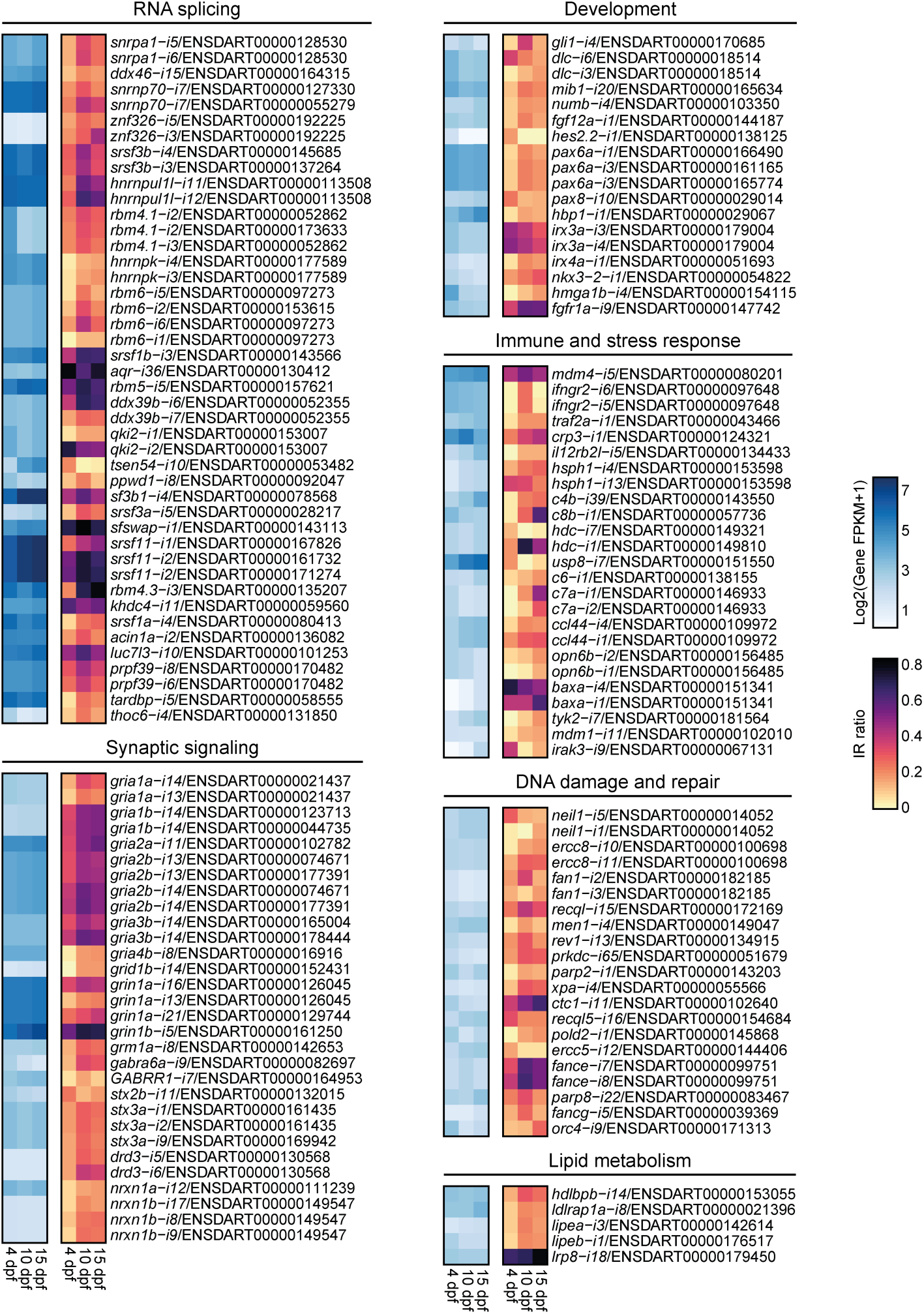
Comparative analysis of transcript abundance and intron retention levels during zebrafish larval development. Heatmaps showing transcript levels (left) and IR ratios (right) for a manually curated set of differentially retained introns. Genes are grouped by functional annotation: RNA splicing, synaptic signaling, development, immune and stress responses, DNA damage and repair, and lipid metabolism. For genes with multiple differentially retained introns, each intron is uniquely labeled with gene name, intron number, and ENSEMBL transcript ID of the representative isoform.

**S3 Fig.**
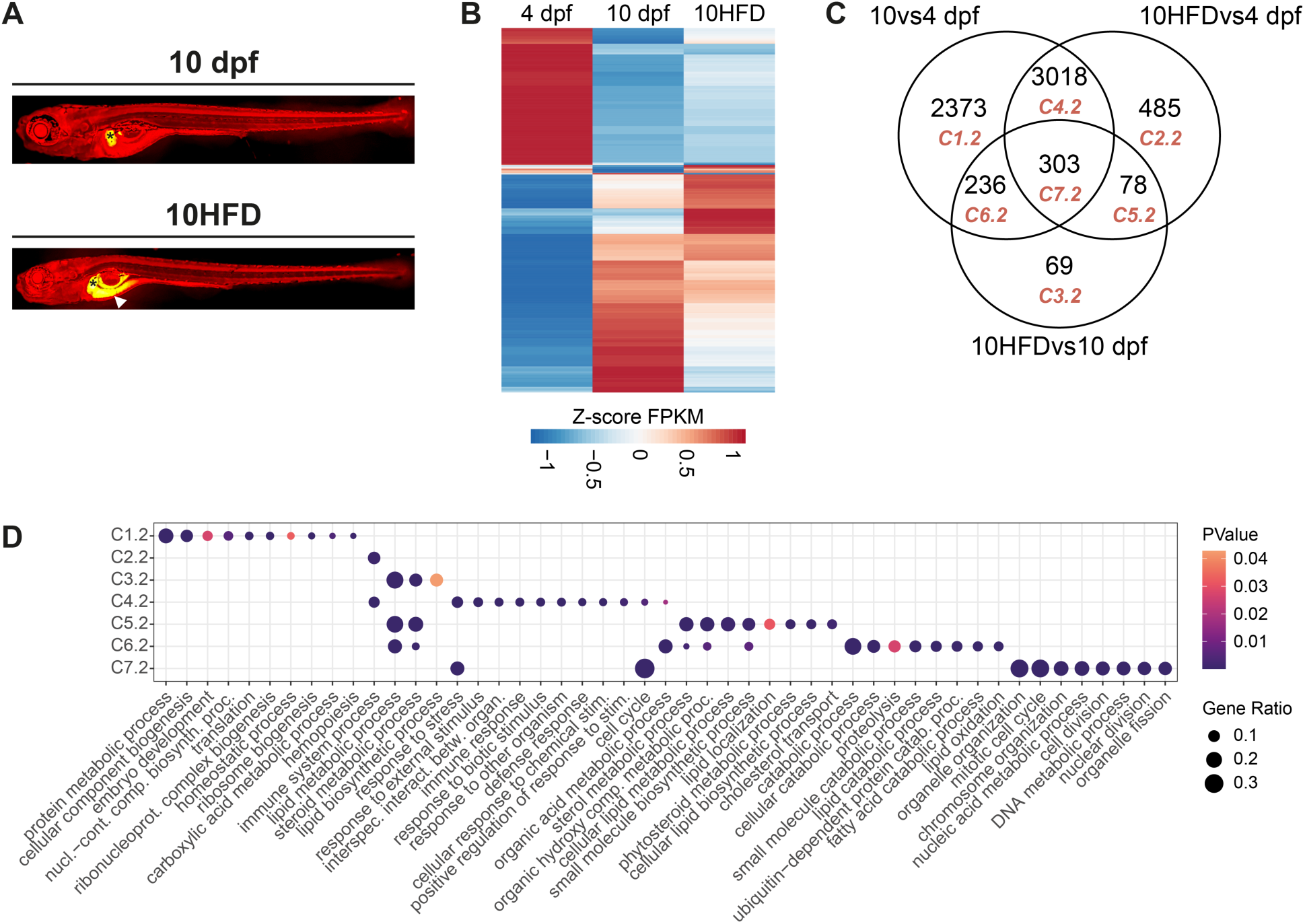
Transcriptomic analysis of zebrafish larvae fed with a high-fat diet (HFD). **(A)** Representative Nile Red-stained images of 10 dpf larvae fed a standard diet (10 dpf) or high-fat diet (10HFD). Asterisks and arrows mark lipid accumulation in liver and intestinal tract, respectively. **(B)** Heatmap of Z-score normalized transcript levels for 6,562 differentially expressed (DE) genes across 4 dpf, 10 dpf, and 10HFD samples. **(C)** Venn diagram showing overlap of DE genes among the three contrasts. **(D)** Dot plot showing significantly enriched GO biological processes for each DE gene set from **(C)**. Gene Ratio indicates the proportion of genes associated with each term.

**S4 Fig.**
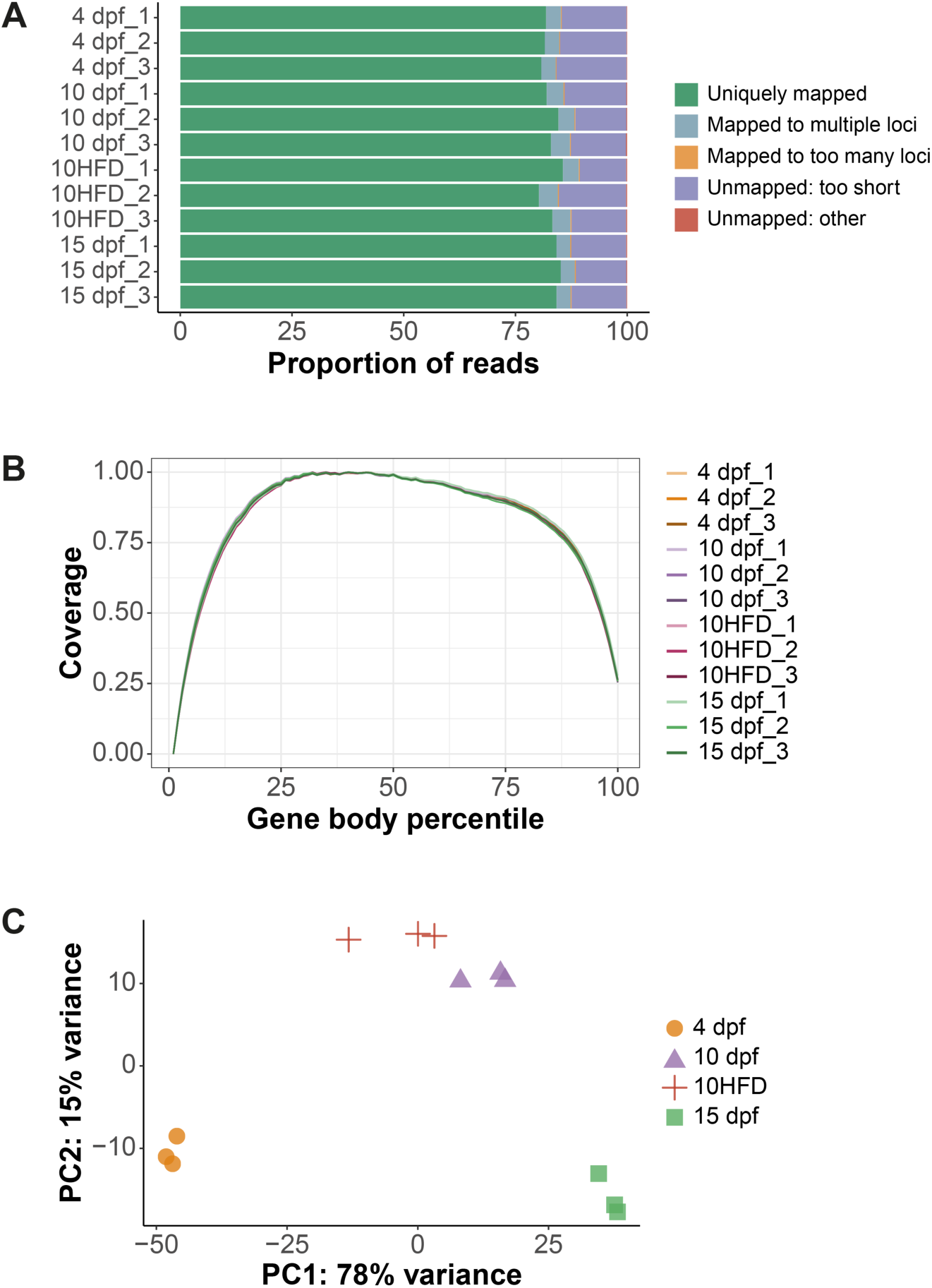
Quality control metrics for RNA-seq data. **(A)** Summary of read mapping quality. The majority of reads aligned uniquely to the reference genome, indicating high mapping specificity. **(B)** Gene body coverage analysis showing uniform read distribution along transcripts. The x-axis represents relative transcript position, with 0 and 1 corresponding to the 5′ and 3′ ends, respectively. **(C)** Principal component analysis (PCA) based on the top 500 most variable genes. Samples cluster by condition and segregate by developmental time point, indicating high biological reproducibility and data quality.

### Supplemental tables

**S1 Table.** Differentially retained introns. Lists of differentially retained introns identified in the contrasts 10 vs 4 dpf, 15 vs 4 dpf, 15 vs 10 dpf, 10HFD vs 4 dpf, and 10HFD vs 10 dpf.

**S2 Table.** SE events. Lists of skipped exon (SE) events identified in the contrasts 10 vs 4 dpf, 15 vs 4 dpf, 15 vs 10 dpf, 10HFD vs 4 dpf, and 10HFD vs 10 dpf.

**S3 Table.** A5SS events. Lists of alternative 5′ splice site (A5SS) events identified in the contrasts 10 vs 4 dpf, 15 vs 4 dpf, 15 vs 10 dpf, 10HFD vs 4 dpf, and 10HFD vs 10 dpf.

**S4 Table.** A3SS events. Lists of alternative 3′ splice site (A3SS) events identified in the contrasts 10 vs 4 dpf, 15 vs 4 dpf, 15 vs 10 dpf, 10HFD vs 4 dpf, and 10HFD vs 10 dpf.

**S5 Table.** MXE events. Lists of mutually exclusive exon (MXE) events identified in the contrasts 10 vs 4 dpf, 15 vs 4 dpf, 15 vs 10 dpf, 10HFD vs 4 dpf, and 10HFD vs 10 dpf.

**S6 Table.** All retained introns. Lists of all retained introns identified in zebrafish larvae at 4 dpf, 10 dpf, 15 dpf, and 10HFD.

**S7 Table.** Differential gene expression analyses. DESeq2 output tables for the contrasts 10 vs 4 dpf, 15 vs 4 dpf, 15 vs 10 dpf, 10HFD vs 4 dpf, and 10HFD vs 10 dpf.

**S8 Table.** Gene set enrichment analysis for differential gene expression (DE). Lists of significantly enriched Hallmark and KEGG gene sets for the differential gene expression contrasts 10 vs 4 dpf, 15 vs 4 dpf, and 15 vs 10 dpf.

**S9 Table.** Gene ontology analysis for DE genes. Lists of GO terms significantly enriched among differentially expressed genes in the contrasts 10 vs 4 dpf, 15 vs 4 dpf, and 15 vs 10 dpf.

**S10 Table.** Gene ontology analysis for genes with differentially retained introns. Lists of GO terms significantly enriched among genes with differential IR in the contrasts 10 vs 4 dpf, 15 vs 4 dpf, and 15 vs 10 dpf.

**S11 Table.** Venn diagram intersections. Lists of differentially retained introns and DE genes corresponding to the intersections displayed in the Venn diagrams presented in Fig 3C and S3C Fig (contrasts 10 vs 4 dpf, 10HFD vs 4 dpf, and 10HFD vs 10 dpf).

**S12 Table.** Gene ontology analysis for genes with differentially retained introns in Venn diagram intersections. Lists of GO terms enriched among genes with differential IR in each intersection from the Venn diagram in Fig 3C (contrasts 10 vs 4 dpf, 10HFD vs 4 dpf, and 10HFD vs 10 dpf).

**S13 Table.** Gene ontology analysis for DE genes in the Venn diagram intersections. Lists of GO terms enriched among DE genes in each intersection from the Venn diagram in S3C Fig (contrasts 10 vs 4 dpf, 10HFD vs 4 dpf, and 10HFD vs 10 dpf).

## Notes

### Competing Interest Statement

The authors have declared no competing interest.

